# Regional Differences in Brain Plasticity and Behaviour as a Function of Sex and Enrichment Type: Oxytocin Matters

**DOI:** 10.1101/2021.05.26.445890

**Authors:** Jamshid Faraji, Hamid Lotfi, Alireza Moharrerie, S. Yaghoob Jafari, Nasrin Soltanpour, Rosa Tamannaiee, Kameran Marjani, Shabnam Roudaki, Farhad Naseri, Reza Moeeini, Gerlinde A.S. Metz

**Author notes:** Authors contributed equally (Jamshid Faraji, Hamid Lotfi, Shabnam Roudaki). **Corresponding author:** Gerlinde A.S. Metz, PhD, Canadian Centre for Behavioural Neuroscience, University of Lethbridge, 4401 University Drive, Lethbridge, Alberta T1K 3M4, Canada.

## Abstract

The early environment is critical to brain development, but the relative contribution of physical vs. social stimulation is unclear. Here, we investigated in male and female rats the response to early physical and social environmental enrichment in relation to oxytocin (OT) and brain-derived neurotrophic factor (BDNF) expression. The findings show that males and females respond differently to prolonged sensorimotor stimulation from postnatal day 21-110 in terms of functional, structural and molecular changes in the hippocampus vs. medial prefrontal cortex (mPFC). Physical enrichment promoted motor and cognitive functions and hippocampal BDNF mRNA and protein expression in both sexes. Combined physical and social enrichment, however, promoted functional and structural gain predominantly in females. These changes were accompanied by elevated plasma oxytocin (OT) levels and BDNF mRNA expression in the mPFC while the hippocampus was not affected. Administration of an OT antagonist in females blocked the beneficial effects of enrichment and led to reduced cortical BDNF signaling. These findings suggest that an OT-based mechanism selectively stimulates a region-specific BDNF response which is dependent on the type of experience.

## Introduction

Environmental enrichment (EE) involving sensorimotor and social stimulation is recognized as one of the most powerful influences on neuronal plasticity (*1-6*). Hence, EE has become a classic experimental paradigm to study the impact of experiences on the brain, and the treatment of psychiatric and neurological diseases (*7, 8*) and stress (*5, 6, 9, 10*). Social interaction represents an integral component of EE and reveals pronounced sex differences (*11, 12*). Females in general respond more prominently to social enrichment than males (*3, 4*), with some exceptions (*13*). Neurohormonal disparities and procedural variables contribute to these sexual dimorphisms (*14, 15*).

The impact of EE is particularly reflected by the hippocampus (HPC) (*16-19*). The HPC, in conjunction with the medial prefrontal cortex (mPFC), contributes to the processing of emotional and social memories (*19-21*), and is intimately involved in movement (*22*) and stress response (*23*). Moreover, the HPC is particularly responsive to social context (*24, 25*) and receptors for oxytocin (OT), a neuropeptide that promotes social behaviours and bonding (*25, 26*), are abundantly expressed by hippocampal neurons (*27, 28*). The pathway linking social support and OT may be involved in stress response regulation (*29*), cortical plasticity induced by sensory stimulation (*30, 31*) and activation of the dopamine reward systems (*32*). Especially in females, social enrichment was shown to increase hypothalamic OT production along with increased brain-derived neurotrophic factor (BDNF) expression in the mPFC (*3, 4, 33*).

BDNF expression responds to both sensorimotor and social enrichment. It is expressed in response to voluntary physical exercise (*34, 35*) and is essential for synaptic plasticity and HPC neurogenesis induced by sensorimotor stimulation (*36, 37*). BDNF was suggested to be the critical factor linking the benefits of social enrichment to neural plasticity and synaptogenesis (*3, 38*). This association was confirmed by studies of social isolation, which reduces circulating OT levels (*39*) and activity of OT-related neural circuits (*40*), along with reduced HPC volume and BDNF expression (*41*). Thus, social stimulation produced by EE paradigms may occur through a close dialogue between the brain’s OT system and BDNF signaling cascades. The differential impacts of sensorimotor vs. social stimulation and their links to OT and BDNF are not yet known, however. Here, we investigated in rats the response to sensorimotor and social enrichment in relation to OT and BDNF expression. We hypothesized that prolonged sensorimotor enrichment from early life to adulthood promotes sex-specific HPC and mPFC development via an OT-BDNF pathway. We anticipated that OT initiates a sexually dimorphic response to social support which then triggers BDNF. The results suggest that an OT-based mechanism selectively stimulates region-specific BDNF expression in the mPFC that is different from the HPC changes.

## Results

### Prolonged Physical Enrichment Promotes Brain Growth and Sensorimotor Performance

#### Body and brain weight

Male and female rats were assigned to Standard Environment (SE), Simple Enriched Environment (sEE) and Complex Enriched Environment (cEE) (n=10-11) (Figure 1**A**-**D**). Body weight among males showed a main effect of Group (*F*_2,29_=54.79, *p*<0.001, η^2^=0.79; *R-M* ANOVA) and a significant interaction between Group and Day (*p*<0.001) where cEE rats weighed less than SE and sEE rats (227<249.97<249.43g, all *p*<0.05, *post-hoc* Tukey HSD). Furthermore, post-mortem brain weight indicated significant group differences (*F*_2,29_=6.70, *p*<0.05) where cEE rats displayed higher brain weight than other groups (1.703>1.645>1.641g, all *p*<0.05, *post-hoc* Tukey, Figure 1**E**). Females did not show significant group differences in terms of body and brain weight (Figure 1**F**). Thus, sensorimotor stimulation promoted the ratio of brain/body weight in males but not females. Also, there was no significant correlation between body and brain weights (all *p*≥0.05, Figure 1**G**&**H**).

**Figure 1:**
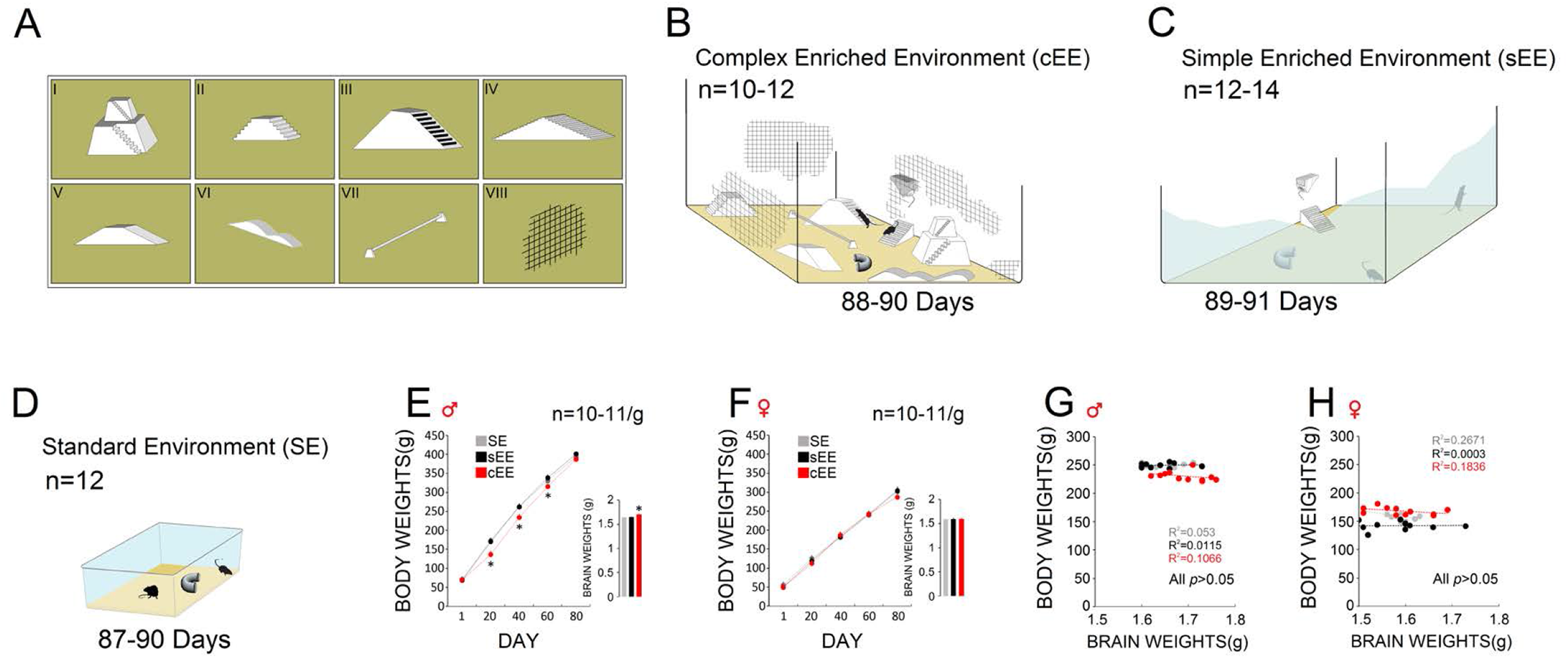
(**A-D**) Male and female rats were raised and housed for 87–91 days in either (**B**) Complex Enriched Environment (cEE) which was equipped with multiple different contraptions (*stepped ziggurat, bilateral smooth and high stairs, bilateral ladder, wavy and smooth slopes, low balance beam*) surrounded by *wire-mesh* covering; (**C**) Simple Enriched Environment (sEE); and (**D**) Standard Environment (SE) units. (**E**) Both body and brain weights in males were significantly impacted by prolonged sensorimotor stimulation. (**F**) Females raised in the cEE condition did not reveal changes in the body and brain weights when compared with SE- and sEE-raised rats. (**G**&**H**) No significant correlation was found between body and brain weights in males and females. Asterisks indicate significant differences: \**p* < 0.05, \*\**p* < 0.01; *One*-*Way* and/or *Repeated*-*measures* ANOVA. Error bars show ± SEM.

#### Sensorimotor function

All animals were able to navigate the BBT (Figure 2**A**). Males (n=9-10) showed a significant effect of Group (*F*_2,25_=19.11, *p*<0.001, η^2^=0.605; *R-M* ANOVA), Trial (*p*<0.001) and interaction between Group and Trial (*p*<0.05) when latency (i.e., the time spent to traverse the bar) on the BBT was considered. cEE rats were able to cross the bar with shorter latency when compared to SE and sEE animals (6.90<10.70<11.84 s, all *p*<0.05, *post-hoc* Tukey HSD). There was no difference between SE and sEE groups (Figure 2**B**). Also, a significant effect of Group (*F* _2,25_=7.92, *p*<0.05, η^2^=0.38; *R-M* ANOVA) indicated that male cEE rats displayed a longer stride length than other groups (4.86>4.11>4.03 cm), particularly on trials 2 and 3 (all *p*<0.05, *post-hoc* Tukey HSD). More importantly, the average stride length in cEE increased across three trials (4.41<4.98<5.18 cm) suggesting that rats gradually gained better balance and performed longer steps. SE and sEE rats did not show these improvements across trials (Figure 2**C**). Housing conditions in females (n=10-11) also resulted in a significant effect of Group (*F*_2,28_=7.71, *p*<0.05, η^2^=0.35; *R-M* ANOVA), Trial (*p*<0.05) and Group by Trial (*p*<0.01) where cEE rats displayed significantly lower latency during beam walking than SE and sEE groups (7.12<9.98>9.77s, all *p*<0.05, *post-hoc* Tukey HSD, Figure 2**D**). Also, female cEE rats showed a larger stride length than SE and sEE groups in trial 2 (3.20>2.78<2.81 cm; *F*_2,28_=4.82, *p*<0.05, η^2^=0.25; *R-M* ANOVA, Figure 2**E**). Thus, sensorimotor stimulation promoted balance and sensorimotor integration in males and females.

**Figure 2:**
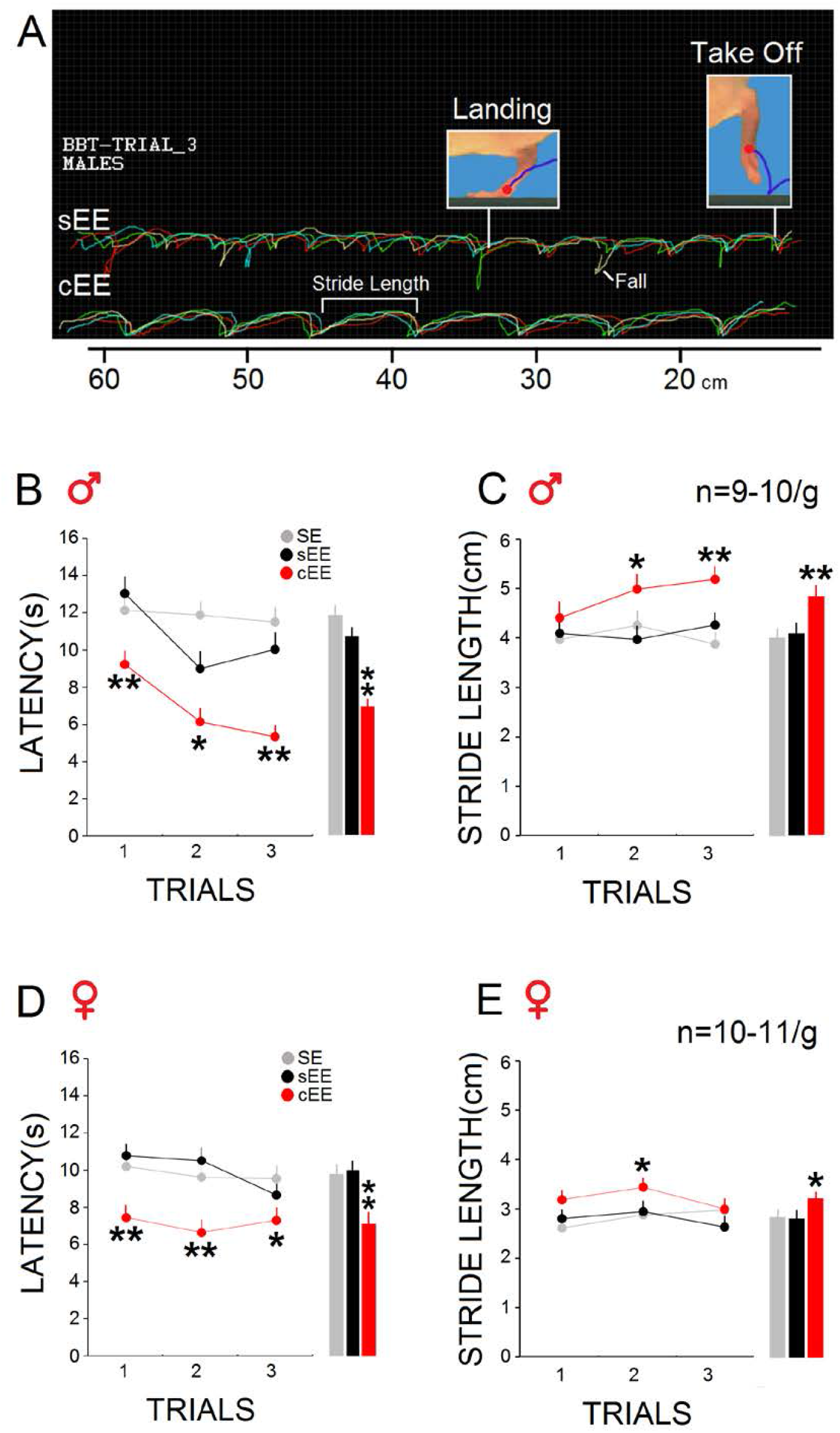
(**A**) Illustration of motion tracks in the balance beam task constructed from four animals in sEE and cEE groups. Each colored motion track represents one animal. Inset pictures display takeoff and landing positions in each stride. Note that stride length, the pixel-based distance between takeoff and landing positions of the left hindlimb, was shorter and more irregular in sEE than in cEE animals. (**B**) **Males**: Latency, the traverse time, indicated a difference between groups across trials where cEE rats were able to cross the bar with shorter latency than other groups. Average latency (s) in all groups is shown in bar graphs. (**C**) cEE animals showed significantly longer stride length on the BBT compared with SE and sEE animals in trials 2 and 3. Average stride length (cm) is shown in bar graphs. (**D**) **Females**: The cEE condition significantly reduced latency on the task in all trials. (**E**) cEE female rats made larger stride lengths than other groups only in trial 2. Average latency (s) and stride length (cm) are shown in bar graphs. Asterisks indicate significant differences: \**p* < 0.05, \*\**p* < 0.01; *One*-*Way* and/or *Repeated*-*measures* ANOVA. Error bars show ± SEM.

### Prolonged Physical Enrichment Facilitates Sex-specific Hippocampal Plasticity and Spatial Learning

#### HPC volume

The HPC volume for each rat was estimated based on 9–10 cross-sections of the hippocampal area (Figure 3**A**). In males (n=7-8) there was no impact of cEE on HPC volume (*p*>0.42, Figure 3**B**). In females (n=6-8) however, cEE housing led to larger HPC volume compared to sEE and SE groups (*F*_2,23_=5.23, *p*<0.05; *O-W* ANOVA, Figure 3**C**).

**Figure 3:**
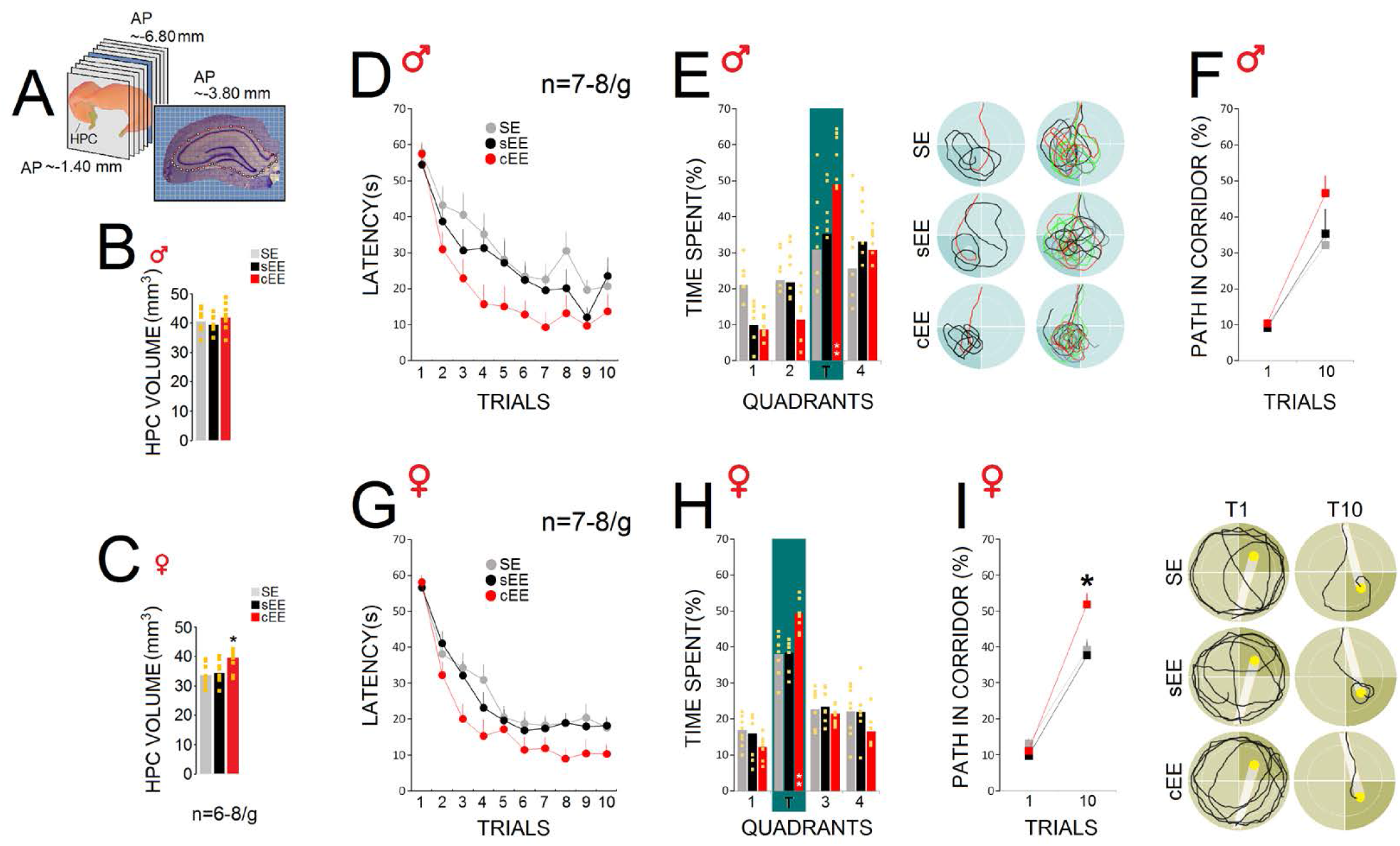
(**A**) A set of 9–10 cross sections of whole hippocampus (AP: −1.40 mm −6.80 mm) that were considered for volumetric analysis along with a coronal view of a right dorsal hippocampus illustrating the area that was considered for hippocampal volumetrics. (**B**) HPC size in males was not impacted by the cEE-housing condition. (**C**) Females raised in cEE showed larger HPC sizes compared to the sEE and SE groups. (**D**) cEE male rats displayed enhanced spatial working and (**E**) reference memory shown by more focal search within the target quadrant (T) of the Morris water maze when compared to sEE and SE groups. (*Right panels*) Representative probe trial trajectories illustrating search patterns of one rat from each group and multipath from four rats within the target quadrant. (**F**) All groups showed similar rates of corridor errors in trials 1 and 10. (**G**) cEE females acquired and retrieved the spatial location of the hidden platform more quickly than sEE and SE groups. (**H**) cEE female rats spent more time than other groups searching in the target quadrant (T) in which the hidden platform had previously been located. (**I**) Analysis of path in corridor (12 cm) for the first and last trials showed that cEE rats swam in the corridor in trial 10 more than other groups. (*Right panels*) Representative swim paths show corridor errors in the first and last trials made by one rat from each group during spatial navigation. Grey strips in plots represent required swim corridor to the platform (Yellow squares in the bar graphs represent individual animals). Asterisks indicate significant differences: \**p* < 0.05, \*\**p* < 0.0; *One*-*Way* and/or *Repeated*-*measures* ANOVA. Error bars show ± SEM.

#### Spatial learning

In male rats (n=7-8), spatial performance in the MWT was tested using a one-day assessment protocol (10 trials/animal). All groups acquired and retrieved the location of the hidden platform in a similar manner (Figure 3**D**). However, cEE animals swam significantly faster than sEE and SE groups during spatial navigation (20.12<28.04<32.20 s; *F*_2,19_=17.65, *p*<0.001, η^2^=0.65; *R-M* ANOVA). A main effect of Trial (*p*<0.001) but no interaction between Group and Trial (*p*=0.92) was observed. Overall, cEE rats located the hidden platform more quickly than other groups. Analysis of swim speed during the 10-trial-acquisition period also showed a relatively flat speed profile across all groups. SE rats navigated more slowly than other groups as testing proceeded. However, no significant changes were observed in swim speed. Furthermore, probe function indicated that cEE rats, in comparison with sEE and SE groups, spent a considerable proportion of their time (48.96>35.27>30.82%; *F*_2,19_=10.82, *p<*0.001; *O-W* ANOVA, Figure 3**E**) searching in the target quadrant (quadrant 3) in which the hidden platform had previously been located. Representative plots of paths are shown in Figure 3**E**-*right panels*. No significant group difference was observed when corridor percent time or swim error in trials 1 and 10 were investigated (Figure 3**F**).

Female rats (n=7-8), regardless of their group identity, were also able to acquire and retrieve the spatial information at a similar rate than males. *R-M* ANOVA, however, showed a significant main effect of Group (*F*_2,20_=17.07, *p*<0.001, η^2^=0.63) suggesting that cEE female rats required less time to find the hidden platform than other groups (19.58<26.21<27.42 s; Figure 3**G**). The main effect of Trial (*p*<0.001) but not Group by Trial interaction (*p*>0.72) was significant. Thus, although all female rats showed a gradual decrease in latency across trials, the cEE group located the platform more quickly than SE and sEE animals. No difference was found in swim speed among groups. Moreover, the probe trial (30-s duration) showed that while all rats spent more time in the target quadrant (quadrant 2), cEE rats still spent more time here than sEE and SE groups (49.41>38.46>38.05%; *F*_2,20_=12.02, *p*<0.001; *O-W* ANOVA, Figure 3**H**). Also, analysis of corridor percent time in the first and last trials (trial 1 and 10) indicated inaccurate swims relative to the platform location in trial 10 only in sEE and SE rats (*F*_2,20_=4.57, *p*<0.05; *O-W* ANOVA, Figure 3**I**). The representative swim paths show corridor errors in the first and last trials made by one rat from each group (Figure 3**I**-*Right panels*). Overall, results in the MWT indicated enhanced working memory in rats that were raised in the cEE-housing condition. Dwell time in the quadrant that formerly contained the platform, indicating enhanced reference memory, was also significantly higher for male and female cEE rats.

### Prolonged Physical Enrichment Promotes Hippocampal BDNF mRNA and Protein Expression

#### BDNF mRNA

Analysis of BDNF mRNA expression in the HPC was based on 3-4 tissue sections per animal and region (∼−1.60 to −4.52 mm relative to bregma, Figure 4**A**). In males (n=4), i*n situ* hybridization revealed that BDNF mRNA in the HPC was abundant and expressed differentially in the CA1, CA3, and DG, but not in the CA2. SE and sEE animals displayed a marked decrease in BDNF mRNA in the CA1 (*F*_2,9_=6.88, *p*<0.01; *O-W* ANOVA), CA2 (*F*_2,9_=24.69, *p*<0.001) and DG (*F*_2,29_=13.61, *p*<0.01). No differences were found in the CA3 (*p*=0.07, Figure 4**B**). The only difference between SE and sEE groups was found in BDNF mRNA expression in the CA2 (*p<*0.05; *post-hoc* Tukey HSD). In females (n=4), BNDF mRNA in all four HPC sub-regions showed a significant increase in cEE compared to the other two groups (CA1: *F*_2,11_=12.89, *p*<0.001; CA2: *F*_2,11_=30.56, *p*<0.001; CA3: *F*_2,11_=8.67, *p*<0.05; *F*_2,11_=22.09, *p*<0.001; *O-W* ANOVA, Figure 4**C**). Thus, prolonged cEE housing had a noticeable impact on mRNA expression in all HPC sub-regions in females.

**Figure 4:**
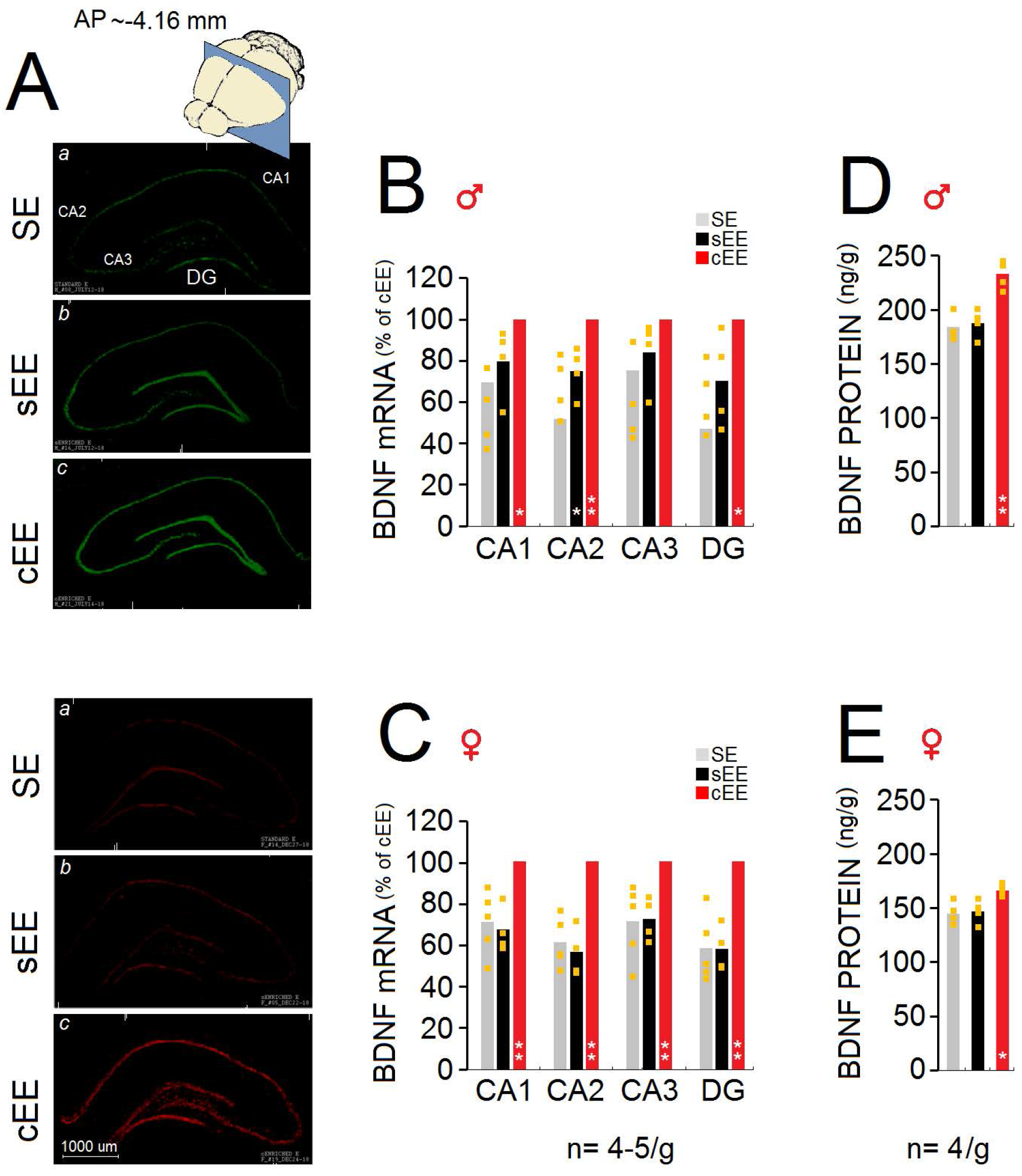
(**A**) Representative autoradiographs (unsharp mask-HK1-RSZII2.60 filter-Green & Red) of the hippocampal BDNF mRNA expression from all groups in males (*top*) and females (*bottom*) illustrate differences in BDNF mRNA expression between groups. Autoradiographs of BDNF mRNA in the dHPC (CA1, CA2, CA3, and DG) indicate that only the cEE housing condition enhanced BDNF mRNA expression in CA1, CA2 and DG in male and female rats. (**B**&**C**) Male and female cEE rats showed the highest BDNF mRNA expression in HPC sub-regions across all groups. (**D**&**E**) Both males and females raised in cEE expressed more BDNF protein in the HPC compared to other groups. (Yellow squares in the bar graphs represent individual animals). Asterisks indicate significant differences: \**p* < 0.05, \*\**p* < 0.01; *One*-*way* ANOVA. Error bars show ± SEM.

#### BDNF protein

BDNF protein expression in males (n=4) revealed no differences between the left and right HPC. However, *O-W* ANOVA indicated group differences in terms of BDNF protein (*F*_2,9_=18.11, *p*<0.001), where rats raised in the cEE condition showed higher BDNF protein expression in the HPC compared to sEE and SE groups (233.25>188>184 ng/g; Figure 4**D**). No difference was observed between sEE and SE groups (*p*=0.90). In females (n=4), the impact of cEE-housing on BDNF protein expression was replicated (*F*_2,9_= 6.56, *p*<0.05) indicating that cEE housing increased BDNF protein expression relatively to sEE and SE conditions (164.75>145.25>143.75 ng/g; Figure 4**E**). Overall, HPC BDNF mRNA and protein expression was raised by cEE housing in both males and females.

### Oxytocin Reflects Synergistic Benefits of Social and Physical Enrichment

#### Body and brain weight

Female rats were housed in groups of 13-14 in csEE and ssEE units for approximately 81 days (Figure 5**A**-**B**). Despite higher body weights in csEE rats compared to ssEE rats in the course of experiment, *R-M* ANOVA did not show significant group differences (*p*=0.058; Figure 5**C**). However, there was a significant group difference in terms of brain weights (*F*_1,25_=4.65, *p*<0.05; *O-W* ANOVA) where the csEE rats displayed higher brain weights than ssEE rats (1.60>1.53g; Figure 5**D**). Body and brain weights showed no correlation (all *p*<0.05; Figure 5**E**).

**Figure 5:**
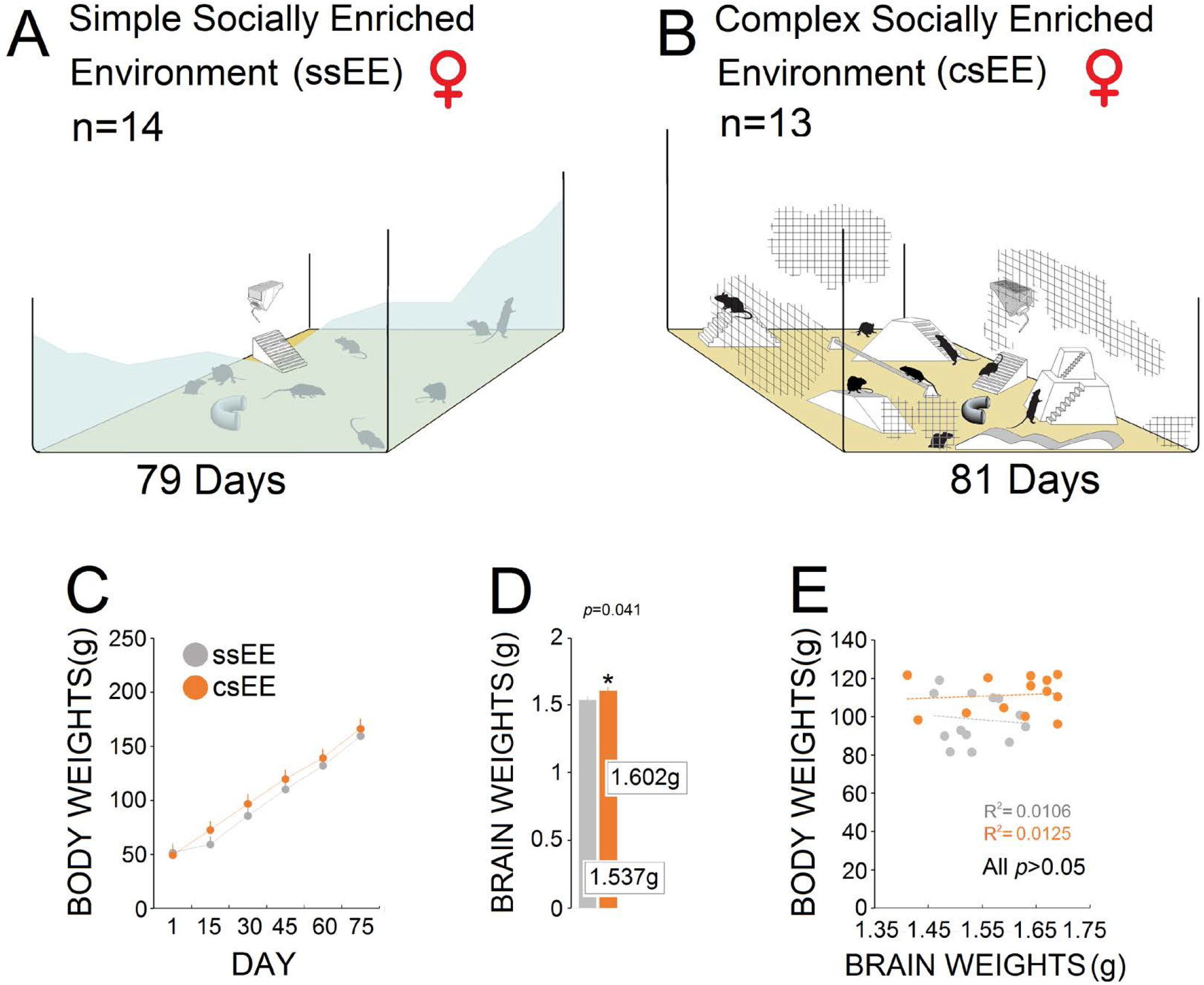
(**A**&**B**) Female pups were housed for approximately 81 days in groups of 14 and 13 in two conditions: (*i*) Simple Socially Enriched Environment (ssEE) and (*ii*) Complex Socially Enriched Environment (csEE) within the environments. (**C**&**D**) Housing condition had no effect on body weight, but csEE did increase brain weight. (**E**) There was no significant correlation between body and brain weights. Asterisk indicate significant differences: \**p* < 0.05, *One*-*Way* ANOVA. Error bars show ± SEM.

#### Brain volume and cortical thickness

The synergy between social experience and sensorimotor stimulation in females caused a trend in larger HPC (*p*=0.361) and brain volumes (*p*=0.054) (n=5-6), with slightly larger brains in csEE compared to ssEE rats. However, csEE rats showed significantly greater volumes in the rostro-caudal sections 2-15, mostly corresponding to the medial prefrontal cortex (mPFC) and primary motor cortex (M1; all *p*<0.05; *O-W* ANOVA; Figure 6**A**). Thus, brain volume was partially impacted by csEE indicating local sensitivity of the brain to the synergy between social experience and sensorimotor stimulation. Furthermore, cortical thickness revealed growth in the medial (*left*: 1.67>1.46 mm, _1,11_=11.03, *p*<0.01; *right*: 1.65>1.48 mm, *F*_1,11_=6.37, *p*<0.05) and lateral (*left*: 1.50>1.30 mm, *F*_1,11_=7.59, *p*<0.05; *right*: 1.55>1.32 mm, *F*_1,11_=13.91, *p*<0.01) portions of the cortex in the csEE compared ssEE rats. The ventrolateral portion in both hemispheres remained unaffected (all *p*<0.05; Figure 6**B**). Hence, cortical thickness confirmed the regional susceptibility of the female brain to social and sensorimotor experiences.

**Figure 6:**
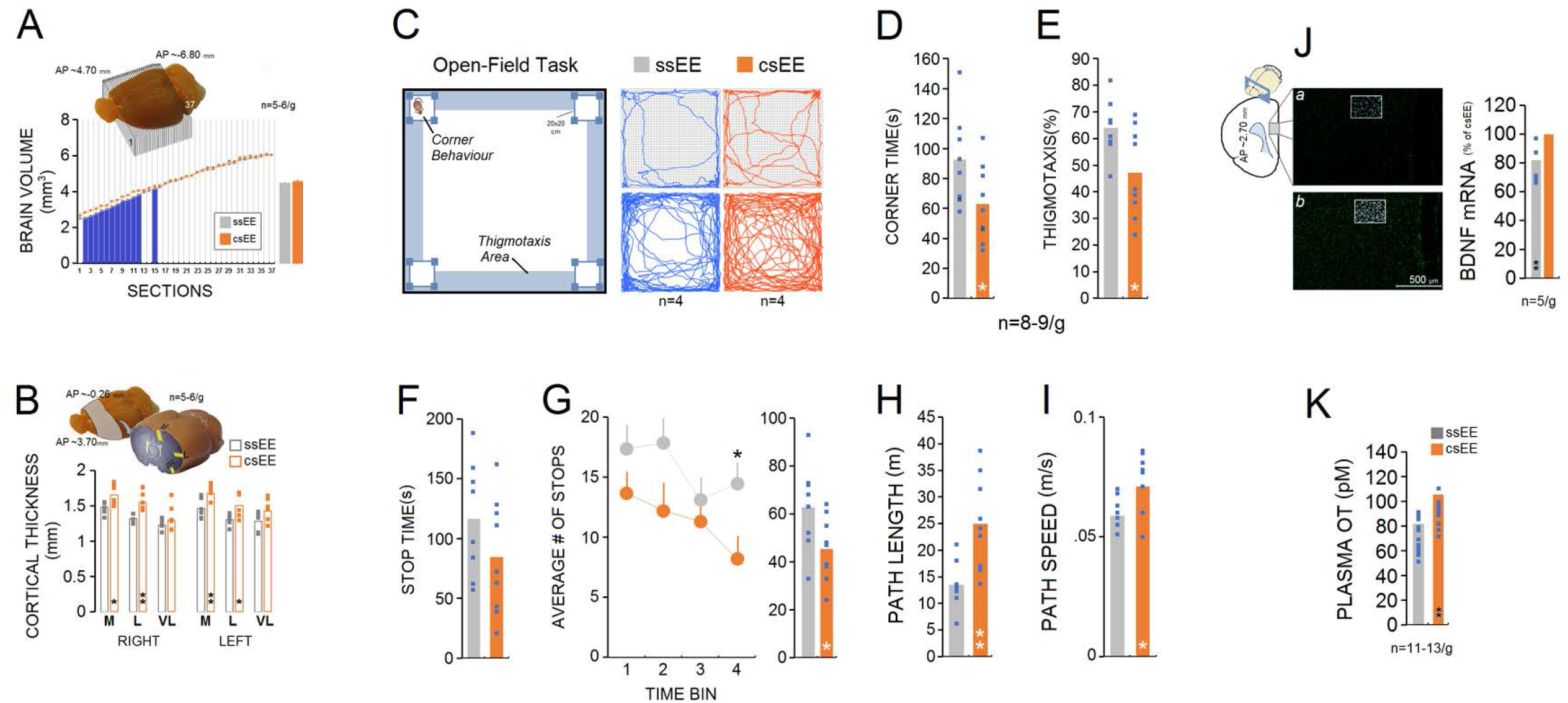
(**A**) Brain volumetric analysis in females considered 36-37 cross sections of the brain except olfactory bulb and cerebellum (n=5-6/g). Brains of the csEE group were larger than those of the ssEE group on the second to fifteenth rostro-caudal brain sections. (**B**) Coronal-sagittal views of the brain illustrate anatomical location of the sections investigated for cortical thickness and the three cortical points (medial, lateral, ventrolateral) used for cortical thickness measurements in both hemispheres. csEE rats showed greater cortical thickness in medial and lateral but ventrolateral portions compared to ssEE rats in both hemispheres. (**C**) Schematic representation of the open field task along with path taken by one representative rat (*top*) from each group and also multipath from four rats (*bottom*). (**D**&**E**) csEE rats spend longer time in the corners and thigmotaxis area than ssEE rats. (**F**&**G**) Despite a similar profile of stop time in both groups, csEE rats made fewer stops than ssEE rats, particularly in the last time bin (the last 2-min). (**H**&**I**) Motivational measures of exploration, such as path length and speed, were influenced by csEE housing. (**J**-*left panel*) mPFC BDNF mRNA signal (scale bar = 500 μm) in the ssEE (*a*) and csEE (*b*) groups. BDNF mRNA data (n = 5/g, *right panel*) are represented as percent of the csEE group where the ssEE rats displayed less mRNA expression in the mPFC when compared with their csEE counterparts (∼21%). (**K**) csEE housing raised plasma OT levels (n=13). Blue columns in panel A represent significant differences between groups. Squares in the bar graphs represent individual animals. Asterisks indicate significant differences: \**p* < 0.05, \*\**p* < 0.01; *One*-*way* ANOVA. Error bars show ± SEM. *M*: medial, *L*: lateral, *VL*: ventrolateral.

#### Exploratory activity

Figure 6**C** depicts the OFT along with paths taken by ssEE and csEE rats. Analysis of corner time (time spent in the corners) and thigmotaxis (repetitive pattern of exploration near to the wall) indicated that csEE rats spent less time in the corners (n=8-9, *F*_1,15_=4.70, *p*<0.05; *O-W* ANOVA) and thigmotactic behaviour (*F*_1,15_=6.01, *p*<0.05; *O-W* ANOVA) when compared with ssEE rats (Figure 6**D**&**E**). Also, both groups displayed similar profiles of stop time in the OFT (*p*=0.189), although the average number of stops in the csEE group was significantly lower than in ssEE rats (*F*_1,15_=5.18, *p*<0.05; *O-W* ANOVA; Figure 6**F**&**G**). Furthermore, path length indicated that the csEE group traveled significantly longer distances than ssEE rats (*F*_1,15_=11.36, *p*<0.01, *O-W* ANOVA). Compared to ssEE animals, csEE rats explored the OFT faster (*F*_1,15_=7.05, *p*<0.05, *O-W* ANOVA; Figure 6**H**&**I**). Thus, social experience associated with sensorimotor stimulation impacted both motivational and emotional aspects of exploratory activity.

#### BDNF mRNA expression

BDNF mRNA levels in the HPC (n=5) were not affected by housing conditions (*dorsal*: *p*=0.40, *ventral*: *p*=0.93). However, the ssEE rats expressed less BDNF signal (∼21%) in the mPFC when compared with the csEE group (*F*_1,8_=13.52, *p*<<0.05; Figure 6**J**). There was no difference between the right and left mPFC (*p*=0.08).

#### Plasma OT levels

Figure 6**K** illustrates changes in plasma OT concentrations in both groups (n=11-13) as a function of housing condition. csEE rats displayed higher levels of circulating OT than ssEE rats (*F*_1,22_=16.38, *p*<0.001; *O-W* ANOVA) suggesting a significant impact of social stimulation on plasma OT (105.48>81.81 pmol/L). There were no correlations between circulating OT concentration and OFT measures.

In summary, prolonged social and sensorimotor experiences are associated with alterations in cortical anatomy, BDNF expression in the mPFC, and circulating OT levels in females.

### Oxytocin Antagonist Prevents Cortical BDNF Expression and Benefits of Enrichment OT

#### antagonist effects in OFT

Figure 7**A**&**B** illustrates the experimental design where one of two groups (n=5-6) raised in csEE units for 81-85 days intermittently received an OT antagonist (L-366,509). OT antagonist treatment significantly impacted locomotion and anxiety-related behaviour (Figure 7**C**-**E**). csEE+OTa rats made more stops particularly in the third (*p*<0.01) and fourth (*p*<0.01) time bins than csEE animals (57.66>37.80; *F*_1,9_=8.27, *p*<0.05; *O-W* ANOVA) along with enhanced thigmotactic behaviours (60.50>37.20%; *F*_1,9_=7.26, *p*<0.05) during exploration in the OFT. Thus, the OT antagonist influenced especially emotional aspects of OFT exploration.

**Figure 7:**
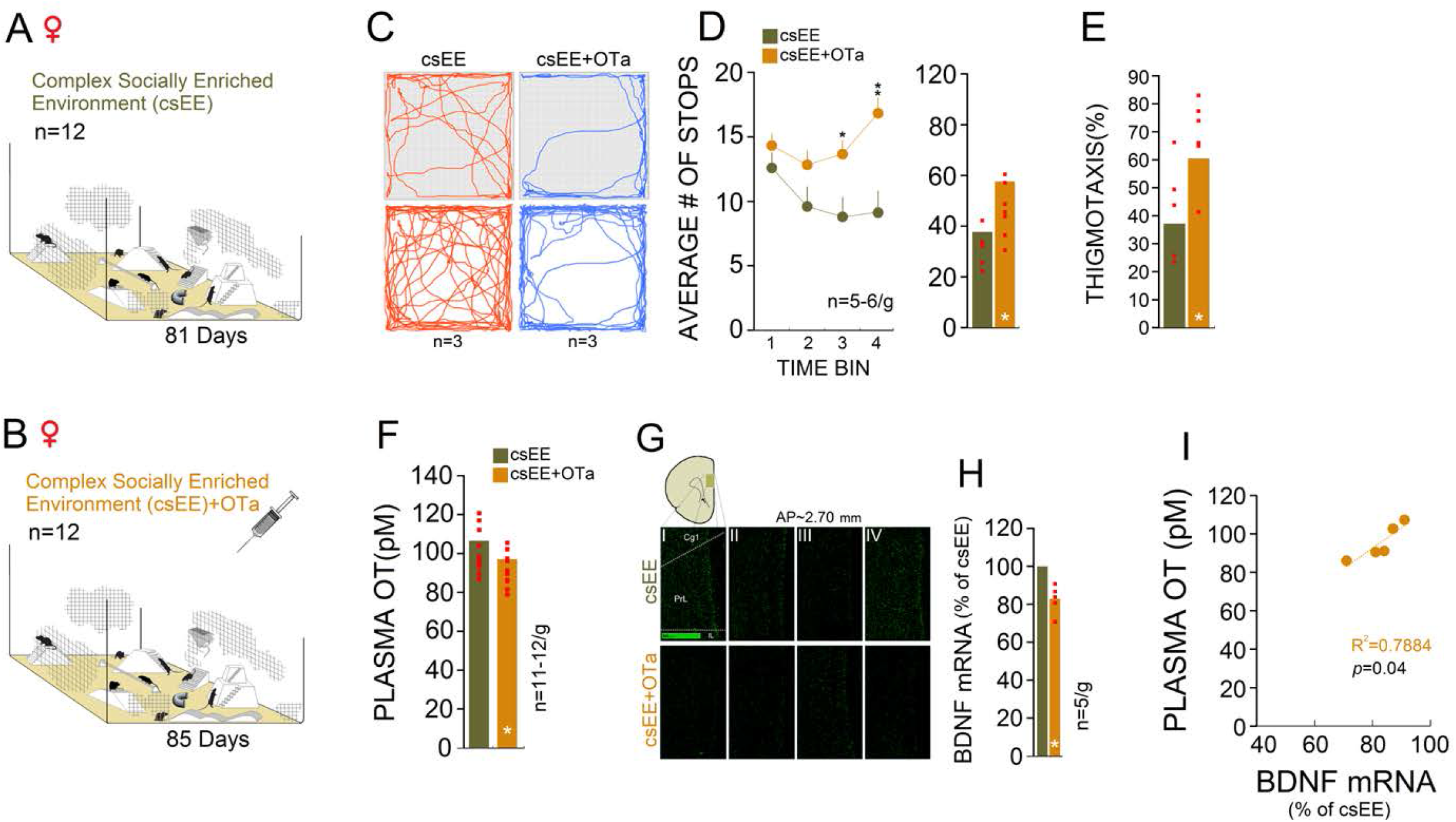
(**A**&**B**) A group of csEE females (csEE+OTa, n=12) intermittently received an OT antagonist (L-366,509). (**C**) Representative exploration paths of one csEE and csEE+OTa rat (*top*) and multipath taken by three rats from each group (*bottom*) in the open field task. Note the differences between paths in the thigmotaxis area in both groups. (**D**) csEE+OTa rats displayed significantly higher number of stops than their csEE counterparts in the third and fourth time bins of an 8-min test session. (**E**) csEE+OTa rats spent more times in the thigmotaxis area when compared with csEE rats. (**F**) OT antagonist administration reduced plasma OT concentration in the csEE+OTa rats. (**G**) Autoradiographs (unsharp mask-HK1-RSZII2.60 filter-Green) represent BDNF mRNA signals in the mPFC in four rats from each group. (**H**) csEE+OTa rats that displayed reduced OT concentration also showed reduced BDNF mRNA expression levels in the mPFC when compared with untreated csEE-only rats. Note that BDNF mRNA data (n= 5/g) are represented as percent of the csEE group. (**I**) csEE+OTa rats showed a significant correlation between plasma OT levels and mPFC BDNF signal. Red squares in the bar graphs represent individual animals. Asterisks indicate significant differences: \**p* < 0.05; *One*-*way* ANOVA. *Cg1*: Cingulate cortex, area 1; *PrL*: Prelimbic cortex; *IL*: Infralimbic cortex.

#### Plasma OT levels

Circulating OT concentration (n=11-12) showed significant differences between groups (*F*_1,21_=4.88, *p*<0.05; *O-W* ANOVA), indicating that the OT antagonist significantly reduced the circulating OT concentration in csEE+OTa rats (106.49>97.06 pmol/L; Figure 7**F**).

#### BDNF mRNA expression

HPC BDNF mRNA expression (n=5) revealed no significant group differences (*dorsal: p*=0.091, *ventral*: *p*=0.063). However, the BDNF mRNA signals in the mPFC in csEE+OTa rats was significantly reduced (∼19%) compared to untreated csEE-only animals (*F*_1,8_=25.86, *p*<0.001; Figure 7**G**&**H**). No significant differences were found between the right and left hemispheres (*p*>0.05). These findings suggest that OT antagonist administration blocked the synergistic effects of prolonged social and sensorimotor experiences on BDNF mRNA signals in the mPFC only. The observed differences were further supported by a significant correlation between OT concentration and BDNF mRNA expression in mPFC of csEE+OTa rats (n=5, *r*=.88, *p*<0.05; Figure 7**I**); similar correlations were not found in the HPC.

## Discussion

The present study demonstrated that prolonged sensorimotor and social stimulation promotes hippocampal-cortical plasticity, cognitive and motor function, in association with BDNF mRNA expression. Both males and females benefited from the cEE condition in terms of sensorimotor integration and spatial learning, with females being slightly more receptive. Moreover, physical enrichment by cEE also raised hippocampal BDNF mRNA and protein expression in both males and females. Females displayed regional susceptibility to combined social and sensorimotor experiences with largest effects in mPFC and motor cortex, which were linked to elevated BDNF expression. Importantly, enhanced mPFC BDNF signals were mediated by OT levels in females in response to the synergy between sensorimotor and social enrichment. These changes, however, were abolished by OT antagonist treatment, suggesting an OT-mediated pathway linking sensorimotor and social context to regional brain plasticity.

Early-life experiences in rodents have a profound impact on motor and cognitive development and maturation (*42-44*) and involve BDNF signaling to shape HPC-mPFC projections in response to psychosocial experiences (*45-48*). The robust projections from the HPC to the mPFC (*49-51*) and the motor cortex (*22, 52*) propose a dynamic process of brain reorganization based on sensorimotor inputs, especially experiences in early life (*53, 54*). Previous insights mainly stem from the classic EE paradigm, which generally involves aerobic exercise (AE) on running wheels in rodents (*19, 55-59*). AE comprises components of aerobic exercise such as cardiovascular stimulation, motivational aspects of behaviour, and general arousal. Reports show that the physiological aspects of AE (e.g. increased cerebral blood volume and blood flow, blood-brain-barrier permeability, angiogenesis and neurogenesis, glucose utilization and increased hormone and growth factor circulation) may explain its robust effects on brain structures such as the HPC and mPFC (*60-62*). By contrast, the present physical EE protocol lacks an apparatus to induce AE, but still encourages musculoskeletal activity through behaviours such as free horizontal and vertical exploration, climbing, and turning.

Although hippocampal interactions with the motor system have been extensively investigated (*22, 52*), hippocampal reliance on sensorimotor inputs during development has received little attention. The present study therefore compared the impact of sensorimotor vs. social context on the maturation of HPC and mPFC and on cognitive vs. sensorimotor functions. Rats typically develop many movement skills by the time they are weaned from their mother at postnatal day 21, but some remain immature by that time (*63, 64*). The present study provided animals with low-intensity sensorimotor stimulation in the cEE condition throughout their adolescence to facilitate ongoing neural plasticity. Interestingly, females displayed more susceptibility to the impact of early sensorimotor stimulation in HPC volume and BDNF expression than males. The cEE paradigm also led to greater accuracy in females’ spatial learning indicated by an increased in-corridor swimming in the last trial. The anatomical measures of brain weight and volume, and cortical thickness in females were most affected by the combination of sensory and social stimulation, however. Interestingly, brain weight in females was not affected by sensorimotor stimulation only. Accordingly, as previously shown by this team (*3*), social stimulation critically influences neuroanatomy and behaviour in females.

Results in the present experiment offer new insights on the dynamic interaction between early sensorimotor and social enrichment that regulates brain structure and function in females via intensified OT action. Previously, we have found that social enrichment is causally associated with increased OT levels and telomere lengths in females (*4*). The neuropeptide OT, which is primarily synthesized in the paraventricular nucleus (PVN) and supraoptic nucleus of the hypothalamus (*65*), mediates early experience-dependent plasticity in the brain (*31*). Moreover, OT not only responds to the activation of sensory circuits (*66*), but also is a key hormonal correlate of social behaviours in mammals (*67, 68*) and regulates brain responses to environmental stimulation in both sexes (*4*). OT may reduce stress and exert anxiolytic consequences particularly when released in response to haptic contact via skin (Uvnas-Moberg et al., 2014). However, a robust effect of sex was reported in OT receptor density throughout the brain (*69*) and in OT receptor signaling in the mPFC of rats (*70, 71*), which may explain why males and females respond differently to social experiences. In the present experiment, the csEE female rats displayed a prominent OT response to the housing condition where sensorimotor and social stimulations were combined for 3 months. Both structural and functional alterations in the presence of increased OT levels disappeared once the endogenous OT was interrupted by an antagonist. This causal, behaviour-to-hormone approach supports the majority of correlational data that link OT to social experiences (*33, 72*). Social cues typically cause OT release in brain regions that are important for triggering or regulating social behaviours. Conversely, the OT may take part in functional changes due to its capacity to modulate emotional and social behaviours. In light of these reciprocal effects which extend to HPC and mPFC, one can hypothesize that the OT system maintains a close dialogue with sensory processing that is critical for detecting and interpreting social cues (*73*). Furthermore, the present findings expand earlier studies in which OT can be released by non-noxious stimulation through stimulation of cutaneous nerves (*74*). Moreover, prolonged sensorimotor and social stimulation in the csEE, as reflected in enhanced BDNF signaling in the mPFC and exploration in the OFT, is arguably also linked to OT-mediated anxiolytic effects (*4, 75, 76*).

The present study suggests that cEE promoted BDNF mRNA signaling in all HPC sub-regions which in turn indicates that both granule cells of the DG and pyramidal cells of areas CA1 and CA3 were receptive to physical activity. Greater susceptibility to the cEE paradigm in females can also be attributed to interactions between 17β-estradiol, BDNF and the HPC mossy fiber pathway (*77*) as it was also shown that 17β-estradiol upregulates mossy fiber BDNF synthesis in female rats (*78*).

Although the signaling pathway linking OT to BDNF regulation is not entirely known, robust OT-induced regulation of BDNF expression and associated neuronal plasticity has been shown in various contexts. OT is able to enhance BDNF protein expression (*79-81*), increase neurogenesis and brain volume (*80*), and modulate behavior via BDNF-TrkB signaling in OT neurons (*82*). Functionally, OT acts via the cAMP/PKA-CREB signaling pathway to gate BDNF actions (*83, 84*). Moreover, OT-enriched mRNAs encode proteins critical for structural and functional plasticity, including cytoskeletal and postsynaptic organization and pathways involved in rapid activity-induced, experience-dependent plasticity (*82*).

## Conclusion

Social context represents a critical determinant of lifetime health and brain function (*4*). The present study compared the impact of sensorimotor vs. social stimulation and for the first time propose a causal link between social stimulation, OT action and subsequent BDNF expression. The data show that early sensorimotor stimulation and social enrichment combined are most powerful in inducing a region-specific response in the mPFC, and that OT is essential to brain maturation and neurodevelopment. The findings suggest combining sensorimotor and social enrichment have synergistic and long-term benefits for brain development and maturation through an OT-BDNF axis. Given that women are particularly vulnerable to various types of neurological and psychiatric disease (*85*), the present findings are relevant for the design of early intervention through physical and social experiences. Understanding the molecular correlates of activity-dependent brain development will advance such approaches to better support lifetime brain health and resilience.

## Materials and Methods

This study involved male and female Wistar rats, bred and raised at the local vivarium. Animals were housed at room temperature (21–24°C) on a 12-hour light/dark cycle (lights on at 7:30 h) with *ad libitum* access to food and water. Body weight was recorded every five days. Prior to behavioural testing rats were handled for approximately 3 min daily for 2-3 consecutive days. All behavioural training and testing was performed during the light phase of the circadian cycle at the same time of day by four male and four female experimenters blind to the experimental groups. All procedures in this study were carried out in accordance with the National Institute of Health Guide to the Care and Use of Laboratory Animals and were approved by the Avicenna Institute of Neuroscience (AINS) Animal Care Committee.

### Experimental Design

*Experiment 1* (Males and Females): After weaning at post-natal day 21 (PND 21), 38 male and 32 female pups gathered from 13 different litters were randomly allocated to one of three housing conditions: (*i*) Standard Environment (SE), (*ii*) Simple Enriched Environment (sEE), and (*iii*) Complex Enriched Environment (cEE). All animals were raised either in SE, sEE or cEE in groups of 2 for approximately 91 days. Once behavioural assessments were completed on day 98-99, all rats were euthanized for morphological assessment. Based on the results and our earlier work which revealed the largest effects of social support in females (*3, 4*), follow-up work in Experiments 2 and 3 focused on female animals.

*Experiment 2* (Females): At weaning (PND 21), 27 female pups were randomly selected. Based on the hypothesis, female pups were randomly split into two groups: (*i*) Simple-Socially Enriched Environment (ssEE) and (*ii*) Complex-Socially Enriched Environment (csEE). Both groups were housed in groups of 14 and 13 animals, respectively, for approximately 81 days. After behavioural testing was completed on day 89, all rats were euthanized for morphological and molecular assessments.

*Experiment 3* (Females): At weaning (PND 21), 24 female pups were randomly selected from 4 different litters and split into two groups: (*i*) Complex-Socially Enriched Environment (csEE; n=12) and (*ii*) Complex-Socially Enriched Environment in which an Oxytocin (OT) antagonist was administered (csEE+OTa). All rats were euthanized on days 91-92 for structural and molecular assessments. For all experiments, only litters consisting of 8–12 pups were used in order to control for the effect of litter size on early dam-offspring relationships. Animals were constantly living in their units until completion of the experiment. Rats in all experiments were randomly assigned to one of the morphological and molecular assays.

### Housing Conditions: Physical and Social Enrichment

*Experiment 1*: (*i*) Standard Environment (SE): Rats assigned to the SE condition were housed and raised in standard sized Polycarbonate cages (43 cm × 29 cm × 19 cm). (*ii*) Simple Enriched Environment (sEE): The sEE unit was a larger Polycarbonate cage (97 cm × 97 cm × 60 cm) with no additional enrichment except for a staircase by which animals were able to reach the water bottle (Figure 1**A-D**). (*iii*) Complex Enriched Environment (cEE): In the large Polycarbonate cage (97 cm × 97 cm × 60 cm), the cEE was equipped with multiple objects wrapped in wire mesh (bird screen) to provide enriched sensorimotor stimulation (Figure 1**A-D**). Males (n=38) and females (n=32) were housed and raised in non-sibling pairs in all three experimental housing conditions.

*Experiment 2*: (*i*) Simple-Socially Enriched Environment (ssEE): Rats in ssEE housing were exposed to a social housing condition in large Polycarbonate cages (97 cm × 97 cm × 60 cm) for 79 days in groups of 14 with no additional enrichment provided. (*ii*) Complex-Socially Enriched Environment (csEE): Rats assigned to csEE housing were raised in groups of 13 for 81 days in large Polycarbonate cages (97 cm × 97 cm × 60 cm) equipped with various objects for sensorimotor stimulation.

*Experiment 3*: To accommodate OT antagonist (OT ANT) administration, animals were split into two complex social housing groups (n = 12/group): (*i*) the csEE, and (*ii*) csEE+OTa where only the latter received an OT antagonist. Animals were housed in large Polycarbonate cages (97 cm × 97 cm × 60 cm) equipped with various objects for 85 days. Both groups were subjected to blood sampling on days 86-87.

EE in the present study on purpose did not include running wheels or other contraptions designed to specifically encourage physical activity. Aspen wood mixed with shredded paper bedding was used in all types of environments and changed once per week.

### Behavioural Assessment

#### Balance beam task (BBT)

The BBT was used to test sensorimotor integration, motor coordination and balance (*86*). Animals were placed on one end of an aluminum square bar (2×2 cm diameter, 130 cm long, and 75 cm high) and their home cage was located at the other end of the bar. A foam pad was placed underneath to cushion a potential fall. The animals were required to cross the bar at least three times, and their movements were video recorded from a lateral view using a digital camcorder (Sony HDR-CX675) at 60 frames/s with an exposure rate of 1 ms. The latency to traverse the bar, the number of times the hind feet slipped off the bar and stride length were recorded. Each stride was defined as the distance between the lift and landing positions of the hind limb. A high-contrast point with proper vertical and horizontal edge definition was chosen on the back of the hind limb (Sony Vegas Pro 11, Japan). Stride length (cm) on the beam was measured by the number of pixels in the tracked frames traced between the lift and landing positions. Also, a suitable target region for tracking was determined based on a pattern that was clearly visible in every frame. If the target point did not contain a high-contrast point to track, the preprocess parameters (e.g. increasing the contrast) were adjusted to make the source image easier to track.

#### Morris Water Task (MWT)

Spatial performance was assessed by a 1-day testing protocol (10 trials per animal) in the hidden platform version of the MWT (155 cm diameter) as previously described (*41*). Briefly, animals were taught to escape from the water (21 ± 1°C) by climbing onto the hidden platform (12 cm diameter). Each trial began with the rat being placed in one quadrant of the pool around the perimeter of the pool in a pseudo-random sequence. The maximum duration of each swim trial was 60 s. The location of the hidden platform remained constant from trial to trial to assess trial-independent spatial learning. A no-platform probe trial was also performed approximately three hours after the completion of the single session hidden platform testing as a measure for reference memory. In the probe trial the platform was removed from the pool and the rats were allowed to swim freely for 30 s. The percentage of time that the animals spent in each quadrant was recorded. The latency to find the hidden platform, swim speed and error index (swim error or corridor percent time) were recorded and analyzed by a tracking system (HVS Image 2020, UK). The error index refers to the accuracy of an animal’s swim trajectory within a 12-cm-wide corridor from the start point to the platform. Any deviation from this corridor during swimming was scored as an error (*87*).

#### Open-Field Task (OFT)

The OFT was used to assess locomotion and anxiety-related behaviour (*9*). The task made of opaque black Plexiglas consisted of a square arena (135×135 cm) surrounded by walls (33 cm height). Each rat was individually placed at the centre of the arena and video recorded under dim illumination for 8 min with a camera mounted above the open field. Video recordings were analysed for path length and path speed, corner behaviour, stop time (speed of 0.0 m/s lasting at least 1 s), and thigmotaxis (percent time spent close to the walls) by the computer tracking system. Stops and thigmotaxis were measured as an indicator of emotion level. Path length and speed in the OFT were considered as indicators of motivation (*9*). The apparatus was cleaned after each animal with 70% alcohol.

### Oxytocin Antagonist Treatment

OT antagonist (OT ANT) was administered to the csEE+OTa group based on earlier descriptions (*4*). The non-peptidyl OT antagonist L-366,509 (MedKoo Biosciences, Inc., Morrisville, USA) (*88, 89*) was administered (60 mg/kg) subcutaneously into the scruff of the neck. Administration occurred every other day (between 11:00 h and 12:00 h noon) for 38–39 days (in total, 38–39 doses/rat) to intermittently inhibit or reduce OT secretion. Administration of L-366,509 started within the first week of the experiment (day 4) and ended 10-13 h before perfusion. The L-366,509 dosage was chosen based on previous reports by this team (*4*) and Kobayashi et al. (*90*). Physiologic saline solution was injected subcutaneously in the csEE group (approximately 37 doses/rat) to control for confounding stress resulting from repeated OT ANT injections.

### Plasma Oxytocin Assay

Blood samples (0.5–0.7 mL) were taken the day after terminating EE housing to measure OT plasma concentration using solid phase radioimmunoassay (RIA) (*4*). All samples were collected in the morning hours between 9:00 h and 11:00 h and no behavioural testing was performed on blood sampling days. Briefly, blood sample tubes containing aprotinin (500 kallikrein inactivation units/mL blood) were centrifuged at 3000 rpm (1700 g) for 15 min at 4°C. Plasma was stored at –70°C until analysis. Sample extraction and processing were performed according to the manufacturer’s manual (Phoenix Pharmaceuticals, Burlingame, CA), and a method previously described by Kobayashi et al. (*90*). Intra- and inter-assay variability was 7% and 15%, respectively, as reported by the manufacturer.

### BDNF Expression and Morphological Analyses

#### In Situ Hybridization

Animals (n= 4-5/group) were euthanized with an overdose of sodium pentobarbital and brains were rapidly removed. All *in situ* hybridization was carried out as previously described [(*41, 91*), with modifications)] and all sections were run under identical experimental conditions. Briefly, brains were sectioned with a cryostat (10-15 µm) approximately from −1.60 to −4.52 mm relative to bregma. The fixed, air-dried sections were incubated overnight with two 33P-labelled 48-mer oligonucleotide probes in hybridization buffer, and excess and unbound probe was washed off. Brain sections were exposed to BioMax MR-1 X-ray films for 25-30 days. Three to four comparable tissue sections per animal and region were considered for further analysis. Levels of BDNF mRNA were analyzed and masked by optical densitometry of autoradiographic films using a computerized image analysis system (MCID, Canada) and ImageJ 1.49p (NIH, USA). BDNF mRNA data in this experiment are represented as percent of cEE group.

#### BDNF Protein Analysis

An enzyme-linked immunosorbent assay (ELISA) was used to quantify BDNF protein levels (*41, 92*). Briefly, the tissues were homogenized in 100 × (w/v) ice-cold homogenization buffer containing a protease-inhibitor cocktail and then diluted to 1:9 in this buffer and then further to a total dilution of 1000×^37^. ELISA was performed on the homogenate using the BDNF Emax Immuno Assay Systems (Promega KK, Tokyo, Japan). The hippocampus (HPC; ∼−3.80 mm relative to bregma) was analysed for BDNF mRNA and protein (n= 3–4 per group) in Experiment 1.

#### Brain Volume and Cortical Thickness Analyses

Procedures for brain volumetry and cortical thickness were adopted and modified from (*93*). <u>Brain volume:</u> Briefly, for each animal, a set of 35-37 cross sections of the whole brain except olfactory bulb and cerebellum stained with cresyl violet was considered for volumetric analysis. Images of the stained sections were captured using an AxioCam (Zeiss, Jena, Germany). The most rostral section measured was located at ∼4.70 mm anterior to bregma and the most caudal section at ∼-6.80 mm posterior to bregma. For each section (R.1x), the contours of the bilateral hemispheres were traced, and their areas were measured using ImageJ 1.47b (NIH, USA). Brain volume averages were calculated by dividing the sum of measures obtained from each brain by the total number of sections (Lost Brain Area in mm^2^). The approximate volume of the brain (Lost Brain Volume in mm^3^) was determined by multiplying the total area in mm^2^ by both the thickness of each slice (40 mm) and the sampling interval (3):

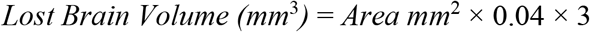

#### Cortical thickness

Three points (medial, lateral and ventrolateral) on 7-8 coronal sections (AP 3.70, 2.70, 2.20, 1.70, 1.60, 1.20, 0.48, and −0.26 mm) from each brain (n=5-6/g) were selected based on Paxinos and Watson (*94*). Therefore, the most rostral section measured was located at ∼3.70 mm anterior to bregma and the most caudal section at ∼ −0.26 mm posterior to bregma. For each point, a vector was considered from the tangent of the outer edge to the inner edge of the cortex. The NDP.view2 viewing software U-12388-01 (Hamamatsu, Japan) was used to record up to six measurements of cortical thickness from each coronal section, three from each hemisphere.

#### Hippocampal (HPC) Volumetry

Three to six hours after behavioural testing, animals were euthanized as described previously (*95*). A series of tissue sections (n= 6–8 per group) were stained with Cresyl violet. HPC volume in each rat was estimated according to the Cavalieri method (*96*) using a set of 9–10 cross-sections of the hippocampal area, starting from −1.40 mm and terminating at −6.80 mm relative to bregma. In the case of missing or damaged sections (less than 9 sections per rat) data were calculated as the average area values from the preceding and following sections.

### Statistical Analysis

The results were subject to analysis of variance (ANOVA; IBM SPSS Statistics, version 21, SPSS Inc., USA). *Repeated measures (R-M), one-way (O-W)* ANOVA, and dependent and independent sample *t*-tests were conducted as necessary. Also, *post-hoc* test (Tukey HSD) was used to adjust for multiple comparisons. In all cases, means of values were compared. Behavioural, morphological and molecular data were analysed by separate ANOVAs with a main between-subject factor of Group. Within-subject factors were designated for each test as appropriate, for example, Body Weight (days 1-80), Trial (1– 10) and Quadrant (target/other quadrants) in the MWT, Trial (1-3) in the BBT, Number of Stops and Time Bins for the OFT, and Regions (HPC and cortex). In order to evaluate the magnitude of the effects of experimental manipulation, effect sizes (*η*^2^ for ANOVA) were calculated. Values of *η*^2^=0.14, 0.06 and 0.01 were considered for large, medium and small effects, respectively. Correlation coefficients were calculated to examine the relationship between OT levels, BDNF expression, etc. In all statistical analyses, a *p*-value of <0.05 (two-tailed) was chosen as the significance level, and results are presented as mean ± standard error.

## Acknowledgements

The authors thank colleagues at the research ethics board at the Avicenna Institute of Neuroscience (AINS) for suggestions and comments, and Dr. M. Fakhamati for his suggestions and assistance with statistical analysis. We gratefully acknowledge the animal care staff at the AINS vivarium for assistance with animal husbandry. We thank S. Saberi, K. Nosrat-abadi, G.R. Farimani and F. Nejad-Ghorban for their assistance with behavioural testing, and N. Schatz for editorial assistance. Funding for this study was provided by a basic science research program-AINS (#41108-010) to RM, and by Discovery Grant #5628 from Natural Sciences and Engineering Research Council of Canada to GM.

## Conflict of Interest

The sponsors had no role in the planning or conducting the study or in the interpretation of the results. The authors declare no potential conflicts of interest with respect to the authorship and/or publication of this article.

## Notes

### Competing Interest Statement

The authors have declared no competing interest.

